# Comprehensive Landscape of Non-muscle Invasive Bladder Cancer Tumour Microenvironment and Prognostic value of Cancer-Associated Myofibroblasts

**DOI:** 10.1101/2023.10.26.564112

**Authors:** Carmen G. Cañizo, Félix Guerrero-Ramos, Mercedes Perez Escavy, Iris Lodewijk, Cristian Suárez-Cabrera, Lucía Morales, Sandra P Nunes, Ester Munera-Maravilla, Carolina Rubio, Rebeca Sánchez, Marta Rodriguez-Izquierdo, Jaime Martínez de Villarreal, Francisco X. Real, Daniel Castellano, Cristina Martín-Arriscado, David Lora Pablos, Alfredo Rodríguez Antolín, Marta Dueñas, Jesús M. Paramio, Victor G. Martínez

## Abstract

**BACKGROUND:** Non-muscle-invasive bladder cancer (NMIBC) poses clinical challenges due to its high recurrence and progression rates. While Bacillus Calmette-Guérin (BCG) remains as the gold standard treatment for high-risk NMIBC, recent irruption of anti-PD-1/PD-L1 drugs claims for a comprehensive understanding of the tumour microenvironment (TME) of these tumors.

**METHODS:** The present prospective study consisted on the analysis 98 fresh NMIBC samples, tumor and non-pathological tissue, via flow cytometry. Final analysis included distribution of 11 cell types and the expression of PD-L1 in 66 tumor and 62 non-pathological tissue biopsies from 73 NMIBC patients (84.4% paired samples). The results were validated using publicly available transcriptomic data, and histology.

**RESULTS:** In comparison to non-pathological tissue, the TME of NMIBC presented microvascular alterations, increased cancer-associated fibroblast (CAF) and myofibroblast (myoCAF) presence, and varied immune cell distribution. Heterogeneous PD-L1 expression was observed across subsets, with cancer cells as primary potential anti-PD-L1 binding targets. Unbiased analysis revealed that myoCAF and M2-like macrophages are enriched in high grade NMIBC tumors, but only myoCAF were associated with higher rates of progression and recurrence, as we confirmed in three independent transcriptomic cohorts (888 total patients). We further validated the prognostic value of myoCAFs by tissue micro-array.

**CONCLUSION:** This comprehensive analysis provides a roadmap to establish the full landscape of the NMIBĆs TME, highlighting myoCAFs as potential prognostic markers.

**FUNDING:** This study was funded by FC AECC (INVES222946GARC), Consejería de Educación, Ciencia y Universidades de la CAM (2018-T2/BMD-10342), Hoffmann-La Roche, Ministerio de Ciencia e Innovación (INMUNOEPIBLA) and ISCIII/FEDER (CIBERONC CB16/12/00489)

## INTRODUCTION

Bladder cancer presents an estimated number of prevalent cases of 491,243 worldwide for both sexes, according to the global cancer observatory for 2022 (https://gco.iarc.fr/en). Of all bladder cancers, approximately 70% are classified as non–muscle-invasive bladder cancer (NMIBC). Patients diagnosed with high risk NMIBC are treated by tranasurethral resection of the tumor followed by immunotherapy with Bacillus Calmette-Guérin (BCG). Although BCG therapy presents a relatively good clinical response, these patients undergo constant invasive surveillance and repeated surgeries due to the high rates of recurrence and progression of these tumors, which elevates the economic cost of this diseases (1) and severely impact the quality of life of the patients (2). Despite many efforts to improve risk stratification of NMIBC patients (3, 4), there is a limited ability to accurately predict which tumors are most likely to recur and/or progress to muscle invasive disease (5). While transcriptomic classification of primary NMIBC tumors has shown good results in terms of prognosis (6–9), these analysis focus mostly on cancer cells, indicating that the study of other components of the tumor may aid in improving these patient stratification methods.

The tumour microenvironment (TME) varies among tumours, main elements including immune cells, stromal cells, blood vessels and non-cellular components such as the extracellular matrix (10). While there is a wealth of works that study the TME in advanced bladder cancer (BLCA) (11–15), this topic has been neglected in non-muscle invasive BLCA (NMIBC), specially for the non-immune compartment of the TME. TME components can impact on therapy response, as has been shown for many solid tumours, including advanced BLCA. Hence, a better characterisation of the TME in NMIBC is important in order to fully understand the biology of these tumours and to improve their management.

The use of anti-programmed death ligand-1 (PD-L1) check point inhibitors is rising as a novel treatment in high risk NMIBC due to limited efficacy of BCG therapy (16), where up to 50-70% of BCG-treated patients will experience a high-grade recurrence despite the instillations (17). Despite promising preliminary results, the response to these check pint inhibitors varies among patients (18, 19). A deeper insight of the molecules targeted by these immunotherapies is pivotal to identify which patients might respond to them. While assessment of PD-L1 expression is valuable for patient stratification in certain cancers, its usefulness in BLCA is a matter of debate (20), especially in NMIBC (21, 22). Besides, not only tumour cells but other TME subsets express PD-L1, which may impact on therapy response and prognosis (23).

We hypothesized that a better characterisation of the TME composition and PD-L1 expression in NMIBC may help understanding the tumour biology and find new prognostic biomarkers and targets. This work is a prospective study in which tumour and non-pathological tissue from 66 NMIBC patients were analysed. While presence of T cell subtypes are well characterised in NMIBC (22, 24), we focused on less studied myeloid and non-immune populations. Here, we provide a reference map of the frequencies of 11 cell subsets in NMIBC and quantified accurately PD-L1 expression in all cell populations. Utilizing computational tools we found that alpha-smooth muscle actin (aSMA)-expressing cancer-associated myofibroblasts (myoCAFs) serve as prognostic biomarkers in these patients, and validated these findings with different techniques and patient cohorts.

## RESULTS

### Clinical and pathological information

We prospectively collected and processed 98 fresh tumour and non-pathological tissue (NPT) samples according to the schematic in figure 1A. After quality control, we included 66 NMIBC tumour samples and 62 NPT biopsies in the final analysis (84.4% matched tumour-NPT ratio). Experimental pipeline for flow cytometry analysis is represented in figure 1B. Among the 66 tumours, 27 were T1 and 39 were Ta, with grade classifications of 15 grade 1, 18 grade 2, and 33 grade 3 (68% high risk). Supplementary Table 1 offers a detailed summary of clinical and histopathological information.

**Figure 1.**
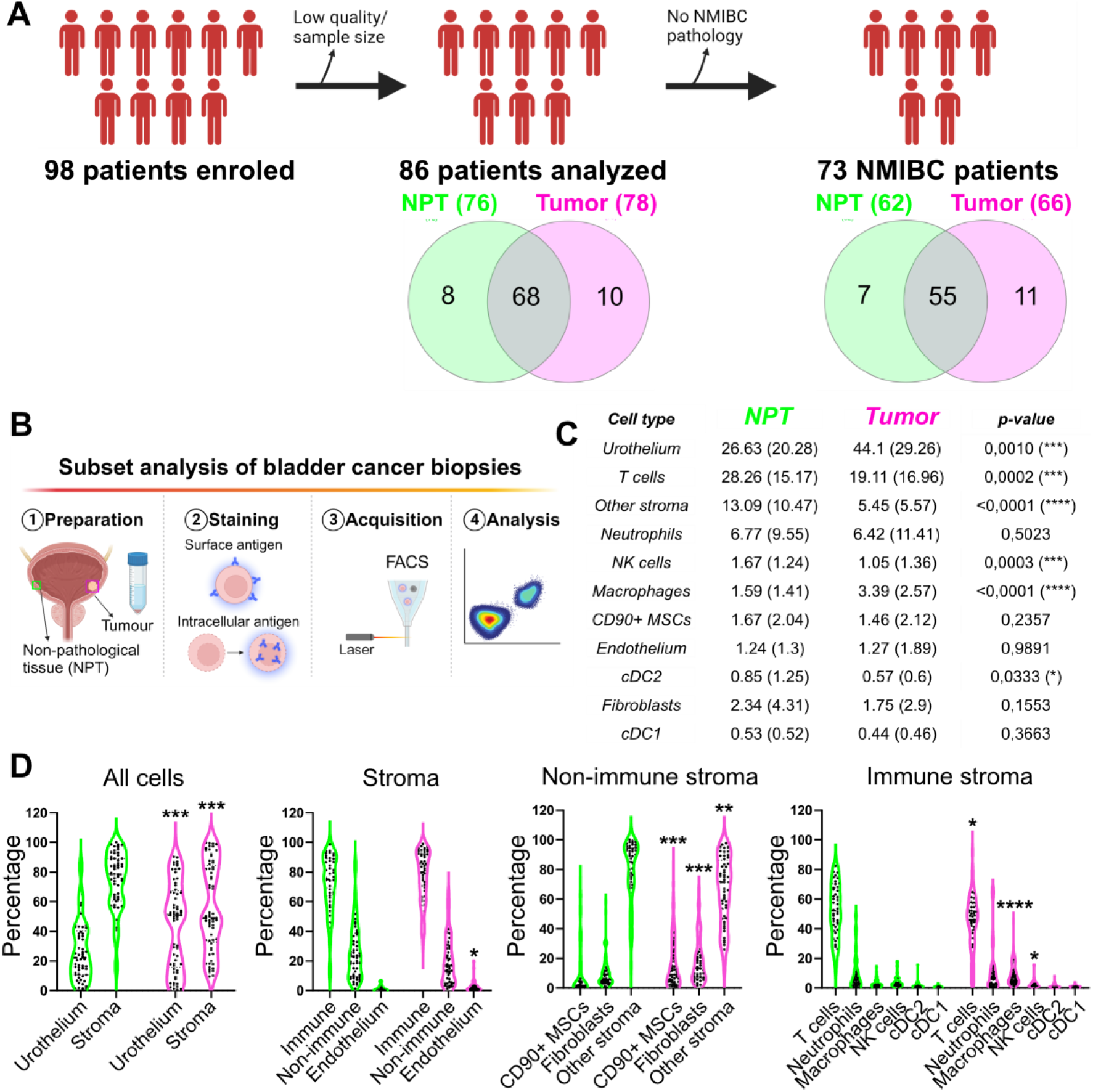
Analysis of cell subset frequencies from total and cellular compartments. A) Patient enrolment and sample analysis. 98 patients were enrolled for the prospective study of which 86 patients were analysed by flow cytometry, excluding samples from 12 patients due to low quality/availability. After pathology analysis, 73 NMIBC patients were included in the final analysis. B) Schematic representation of the prospective study design. Non-pathological and tumour tissue biopsies were collected and digested to obtain single cell suspensions. Samples were then stained for surface and intracellular markers and analysed in a flow cytometer. B) Frequencies for the indicated cell lines were determined for 62 NPT and 66 tumour samples. Mean and standard deviation (in brackets) of the frequencies from total cells for each cell subset is shown. C) Frequencies of cell subsets within the corresponding cell compartment. Each dot represents one patient data. Statistical analysis in B and C was done by Wilcoxon–Mann– Whitney test. * p-value<0.05; ** p-value<0.005; *** p-value<0.0005;**** p-value<0.0001. MSC= mesenchymal stromal cells; NK cells=natural killer cells; cDC=conventional dendritic cells.

### Tumour microenvironment landscape of NMIBC

We designed two multicolour panels for immune and non-immune subset detection, including PD-L1 expression, which allowed identification of 11 cell types (figure 1C). Gating strategy is described in supplementary figure 1A-B. Relative proportions varied widely between NPT and tumours (figure 1C). NPT samples had abundant T cells (28.26 ± 15.17%). Tumours primarily had cancer cells (44.1 ± 29.5%) followed by T cells (19.11 ± 16.96%). Stromal cells, T cells, NK cells, and cDC2 were all decreased in tumors, while macrophages were increased, in accordance to literature (25). To mitigate the impact of the urothelial cell content in tumour samples, we examined cellular compartments separately (figure 1C). TME showed increased endothelial cells, suggesting angiogenesis. All non-immune stroma subsets differed significantly between NPT and tumours. The immune compartment changes paralleled those from whole tissue results, indicating differential recruitment in tumours. We found also fibroblast activation and M2-like differentiation of macrophages increased in tumours (supplementary figure 1C). These results suggest overall cancer-associated inflammation in NMIBC, as expected.

To analyse how the NMIBC TME changes as tumour progresses, we included pT2G3 tumours (n=8) in the analysis (supplementary figure 2). Stromal infiltration increased from pTa to pT2, but leukocyte enrichment remained unchanged (supplementary figure 2A-B and D-E). Only CD90+ Mesenchymal Stromal Cells (MSCs) and Natural Killer (NK) cells showed significant association to pT stage with higher proportion of both cells types in T2 tumors compared to Ta (supplementary figure 2C). Regarding tumour grade, cDC2 were decreased, while CD90+ MSCs and CAFs increased in G3 tumours versus G1 tumors (supplementary figure 2F). Our results reveal that despite the lack of muscle invasion, NMIBC tumours present a diverse microenvironment with prominent recruitment of structural and myeloid cells.

### Increased PD-L1 expression in NMIBC comes from the TME

Anti-PD-L1 therapies have emerged for NMIBC management, prompting PD-L1 expression assessment in tumour compartments. NMIBC presented elevated bulk PD-L1+ cells vs. NPT (figure 2A & B, upper panels). Notably, increase in PD-L1+ cells in tumours stemmed mainly from both immune and non-immune infiltrate, since cancer cells showed similar PD-L1+ proportion to urothelium (figure 2A & B). Subset analysis shows that macrophages had the most PD-L1+ cells, with anti-inflammatory M2-like macrophages especially rich in PD-L1 (figure 2C). cDC2 and tumour cells followed, the later averaging 20% (± 14.42) PD-L1+ cells, along with non-immune stromal cells. Of note, aSMA-expressing myoCAFs had higher proportion of PD-L1+ cells than total CAFs. NK and T cells had lower PD-L1 levels as expected. Next, we sought to highlight key anti-PD-L1 treatment potential binding targets in NMIBC. We gated out total PD-L1+ cells and then applied our previous gating strategy (supplementary figure 1A and B). Cancer cells emerged as the primary binding target, constituting 60% of PD-L1+ cells, surpassing macrophages and aSMA+ myoCAFs which showed high proportion of PD-L1+ cells (figure 2D). Notably, cells with less understood PD-L1 function like Neutrophils and T cells accounted together for a relevant 10% of anti-PD-L1 targets. This underscores the need for further investigating PD-L1’s role in TME subsets. While we quantified PD-L1+ cell dynamics with tumour progression in major compartments, no significant correlations emerged between cancer stage or grade and PD-L1+ cell proportions (supplementary figure 3).

**Figure 2.**
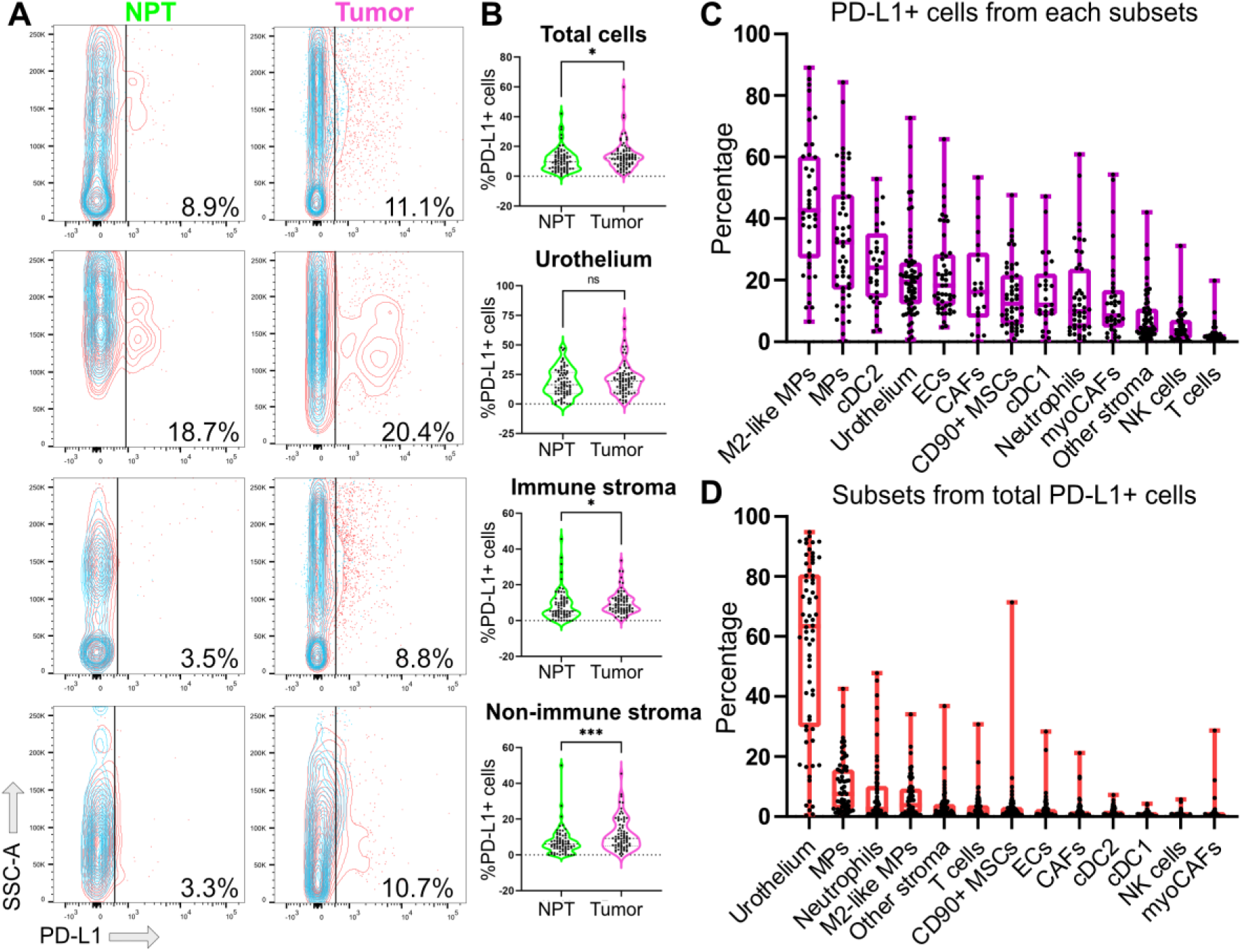
Heterogeneous expression of PD-L1 in cancer and TME cells in NMIBC. A-B) Percentage of PD-L1+ cells was calculated using fluorescent minus one controls (blue overlay) for all cells and main cellular compartments, and representative examples (A) and dispersion plots (B) are shown. Each dot represents one patient data. Comparisons were done by Wilcoxon–Mann–Whitney test. * p-value<0.05; *** p-value<0.0005; ns non-significant. SSC-A= scatter signal channel area. NPT, non-pathological tissue. C) Box plots showing percentage of PD-L1 positive cells within the indicated cell subsets. Each dot represents one different patient. Median and interquartiles are shown. Subsets are ordered from higher median of PD-L1+ cells to lowest. D) PD-L1 positive cells were gated from total cells and subset analysis was run in this set of cells. Each dot represents one different patient. Median and interquartiles are shown. Subsets are ordered from higher to lower frequency median. MP = macrophages; cDC = conventional dendritic cells; ECs = endothelial cells; CAFs = cancer-associated fibroblasts; MSCs = mesenchymal stromal cells; myoCAFs = cancer-associated myofibroblasts.

### M2-like macrophages and myoCAFs are associated to high grade NMIBC

We used our flow cytometry data to identify markers linked to aggressive NMIBC. Employing computational tools in an agnostic approach, we first fine-tuned our method by comparing NPT and tumour samples, verifying findings obtained with the unbiased approach paralleled those from the manual gating (supplementary figure 1 and 4). Subsequently, we focused on tumour samples and generated optimized clustering in both datasets (figure 3A and F), revealing multiple high-grade enriched cell clusters. In the myeloid panel, cluster k7-5 exhibited macrophage markers and variable M2-like-associated markers (figure 3B-D). Manual gating confirmed the enrichment of total macrophages and M2-like macrophages in high-grade NMIBC, aligning with prior research (26). For the stromal/lymphoid dataset, two high-grade enriched clusters emerged, although only one could be defined with the available markers (figure 3G-I). Cluster k15-04 presented high CD90 and podoplanin expression, hence being designated as cancer-associated fibroblasts (CAFs), and showed expression of aSMA. Manual gating confirmed the enrichement of total CAFs and aSMA+ myoCAFs in high-grade NMIBC (figure 3E and J). In summary, our unbiased analysis pinpointed two cell types with distinct functional phenotypes enriched in high-grade NMIBC, holding promise as potential cellular prognostic biomarkers.

**Figure 3.**
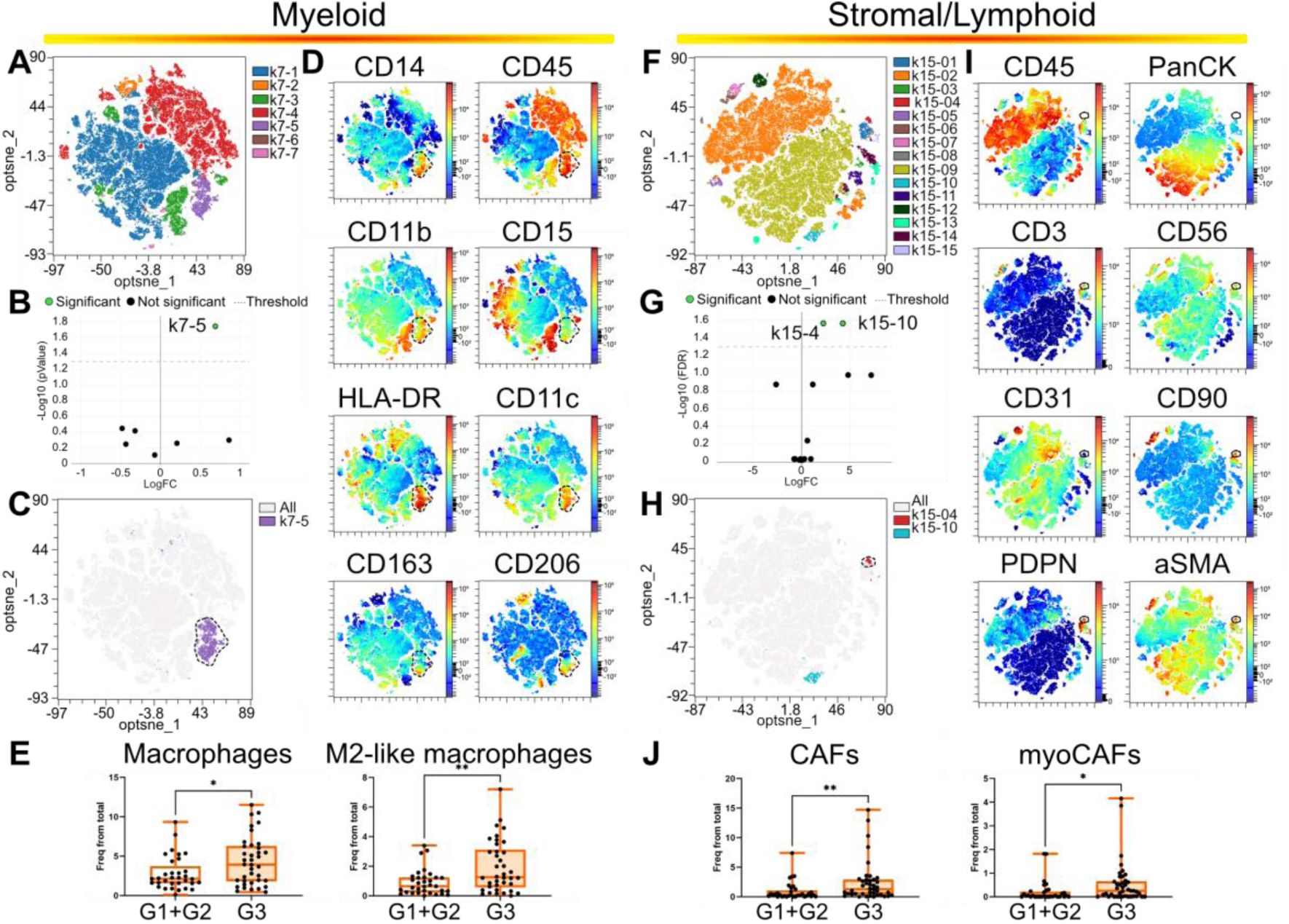
Total and specific macrophage and fibroblast subsets are enriched in high grade NMIBC. Computational analysis of flow cytometry data from the myeloid panel (A-E) and stromal/lymphoid panel (F-J). (A and F) Dimensions reduction was done using optSNE and semi-supervised clustering by FlowSOM setting at 7 and 15 the number of metaclusters for the myeloid and stromal/lymphoid panel respectively. (B, G) G1/G2 versus G3 comparison was done by EdgeR method to determined clusters underrepresented/enriched in G3 tumours. (C, H) Differentially represented cell clusters are shown in the optSNE maps. (D, I) Color-coded optSNE maps show the expression of the indicated markers. Dotted lines highlight differentially represented clusters. (E, J) Results from manual gating validation for the indicated cell subsets. Each dot represents one different patient. Median and interquartiles are shown. Statistical analysis was done by Wilcoxon–Mann–Whitney test. * p-value<0.05; ** p-value<0.005.

### M2-like macrophages and myoCAF crosstalk changes with tumour progression

Reports show macrophage-fibroblast crosstalk in solid tumours (27), but it is unexplored in NMIBC. We observed a positive correlation between M2-like macrophages and CAFs in NMIBC (figure 4A), suggesting crosstalk. Of note, this correlation decreased in high-grade tumours (figure 4B). To investigate this, we used aSMA and CD163 as markers for CAFs and M2-like macrophages, respectively, in tumour sections. Image quantification hinted at more CAFs in G3 tumours, while we could not confirm higher percentage of CD163+ macrophages (figure 4C). Analysing their spatial relationship, CD163+ macrophages were farther from CAFs in G1 tumours (figure 4D and E). This changed drastically in G2 tumours with closer proximity, while G3 tumours presented increased separation, indicating evolving microenvironment dynamics.

**Figure 4.**
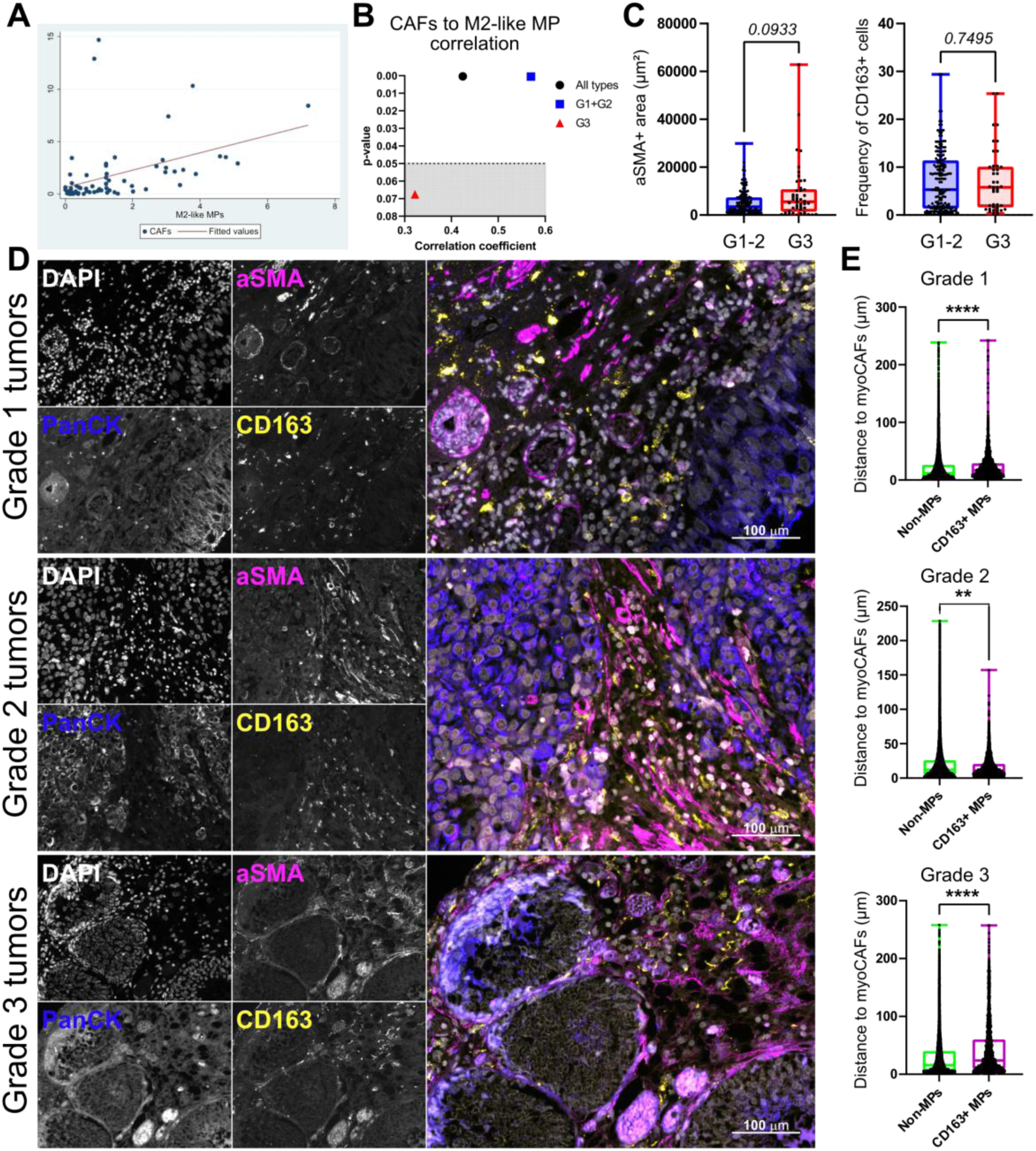
CAF-macrophage interactions decrease in high-grade NMIBC. A) Correlation plot of M2-like macrophage (MP) and CAF frequencies from total cells in all NMIBC samples analysed. B) p-value against correlation coefficient in all samples and G1-2 and G3 groups. Dotted line marks statistical significance for data points above (grey area non-significant). C-E) FFPE tissue sections were stained for Cytokeratins (PanCK), M2-like macrophages (CD163) and myoCAFs (aSMA), plus DAPI to stain nuclei. Using the Qupath software, cytometric analysis was applied to determine the area of aSMA staining, excluding the vasculature, and percentage of CD163 positive cells (C). D) Representative examples for grade 1, 2 and 3 tumour stainings. E) Quantification of the minimum distance to non-vasculature aSMA+ areas for all CD163-negative (non-MPs) and CD163-positive (CD163+ MPs) cells. P-values and significance (asterisks) by Wilcoxon–Mann–Whitney test are shown. ** p-value<0.005; **** p-value<0.0001.

### Macrophage subset gene signatures fail to predict NMIBC prognosis

Given the short follow-up periods of our prospective cohort, we tested the prediction value of macrophages and CAFs as cellular biomarkers by exploring transcriptomic data from NMIBC patients. First, we used single-cell RNA-seq data (14) to identify macrophage clusters. Out of 8 well differentiated clusters, clusters 1, 2, and 3 showed M2 marker expression, thus resembling the M2-like macrophages identified in our flow cytometry data (supplementary figure 5A-B). These clusters also exhibited high scores for immunosuppressive macrophage subsets (28) (supplementary figure 5C). We generated gene signatures for all relevant macrophage subsets (supplementary table 3) and challenged one of the largest NMIBC transcriptomic cohorts, the UROMOL cohort (530 NMIBC patients) (7). All comparisons but one resulted in no association of macrophage gene signatures and recurrence/progression-free survival (supplementary figure 5D-E). We found only a significant association between macrophage cluster 2 and progression-free survival, with patients performing worse when showing low scores for this subset. Overall, our results suggest that immune suppressive macrophage subsets are unlikely to be a significant prognostic factor in NMIBC.

### MyoCAFs serve as cellular predictors of bad prognosis in NMIBC

Next, we assessed the patient cohort with gene signatures generated for total CAFs (panCAFs), inflammatory CAFs (iCAFs), and aSMA-expressing CAFs (myoCAFs), described in various solid tumours, including bladder cancer (12, 14). PanCAFs and iCAFs showed no prognostic value in NMIBC patients for recurrence-free (RFS) and progression-free survival (PFS), confirmed with a well-established panCAF signature (supplementary figure 6A-B). Conversely, the myoCAF signature showed strong association with worse PFS and RFS, with statistical significance for worse performance in patients classified with high scores for myoCAFs (figure 5A). We confirmed this association using an alternative myoCAF signature in the same cohort (supplementary figure 6C). Furthermore, we validated this association in two independent cohorts (6, 9), highlighting a robust link between high abundance of myoCAFs and bad prognosis in terms of recurrence and progression (figure 5A). Furthermore, we noticed that the myoCAF signature was enriched in those patients stratified into the most aggressive transcriptomic classes in all three cohorts (figure 5B). On the other side, panCAFs and iCAFs also associated with the aggressive transcriptomic classes 2a and 2b in the UROMOL cohort (supplementary figure 6D). Nonetheless, when we compared the gene signature scores linked to genomic classes, we found only myoCAF to be highly enriched in the two groups associated to worse RFS and PFS (supplementary figure 6E). These results would indicate a link between the biology of the tumor and the composition of the TME, in this case specifically associated to myoCAF. Finally, explored whether the predictive value of myoCAFs depended on sex. We found similar myoCAF scores in both male and females in two independent cohorts, plus similar predictive value when separating both sexes (supplementary figure 6), concluding that the value of myoCAFs as prognostic factor is independent of sex.

**Figure 5.**
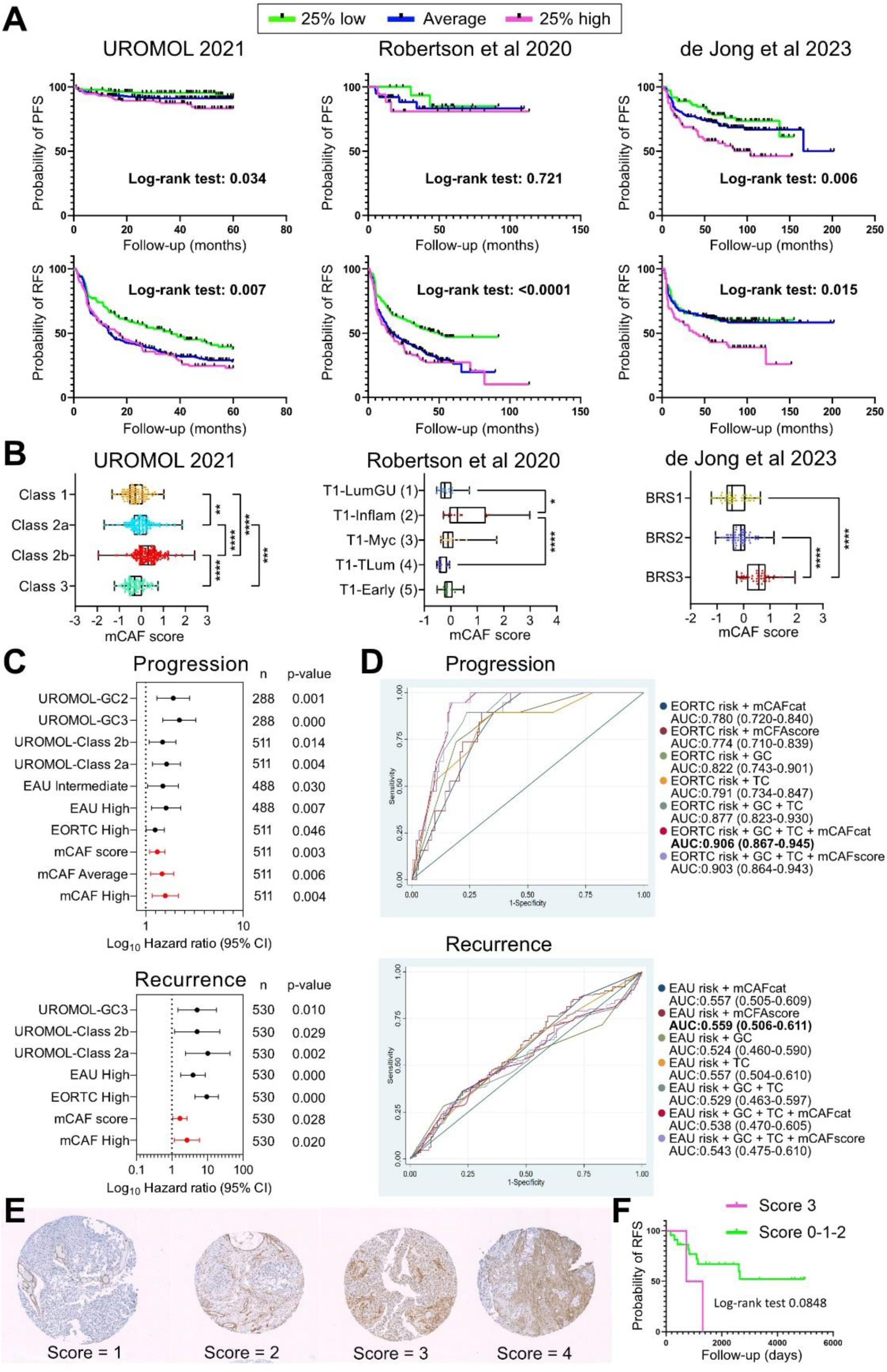
Abundance of myoCAFs associates with bad prognosis in NMIBC. A) A gene signature score was generated for myoCAFs to challenge two independent transcriptomic data cohorts of NMIBC. Patients were ranked according to this score and three groups were formed. Kaplan– Meier plots for probability of progression-free survival (PFS, upper panel) and recurrence-free survival (RFS, lower panels) for the three independent cohorts are shown. Log-rank (Mantel-Cox) test was used to calculate statistical significance between curves. B) Dotplots showing myoCAF scores for patients grouped according to cohort-specific transcriptomic classes. Each dot represents one patient. P-values ** < 0.005; *** < 0.0005; **** < 0.0001. Statistical analysis was done with Kruskal-Wallis test with Dunńs correction for multiple comparisons. D-E) Overview of hazard ratios calculated from univariate Cox regressions of progression-free and recurrence-free survival using clinical and molecular features. Dots indicate hazard ratios and horizontal lines show 95% confidence intervals (CI). P-values and sample sizes, n, used to derive statistics are indicated. EORTC= European Organisation for Research and Treatment of Cancer, EAU= European Association of Urology. D) Performance of a multivariable analysis to predict progression (upper panel) and recurrence (lower panel), considering all the risk factors that had a significant result in the univariate analysis, in the UROMOL 2021 cohort, as well as those that had some relevance among the study results. A step by step selection process was used to highlight the most relevant factors. Receiver operating characteristic (ROC) curves for predicting progression within 5 years using logistic regression models. For each curve, AUC (area under the curve) and confidence interval (in brackets) is shown. Bold letters highlight the model with the highest AUC. FU, follow-up; mCAF-s, myoCAF score; mCAF-c, myoCAF category; mCAF-H, myoCAF category high; Class 2a/C2a, transcriptomic class 2a.F) Representative immunohistochemistry images showing aSMA expression in primary tumour sections from NMIBC patients. Histological analyses and staining was performed in a TMA with duplicates for each tumour (n = 29). All sections were scored from 0 to 3 according to aSMA staining density in non-vasculature areas, to exclude pericytes. G) Kaplan–Meier plot for recurrence-free survival in patients stratified by aSMA staining score. Log-rank (Mantel-Cox) test was used to calculate statistical significance between curves.

Univariate Cox regression identified that both myoCAF category and score strongly predicted PFS and RFS in NMIBC using the UROMOL cohort (figure 5C), with hazard ratios comparable to those of the well stablished predictors EAU and EORTC risk scores in the case of progression. Receiver operating characteristic (ROC) analysis for progression prediction using logistic regression models demonstrated that adding the myoCAF category high improved the prediction power the EORTC risk score, genomic class and transcriptomic class combined together, reaching an outstanding discrimination (AUC: 0.906 ± 0,037) (figure 5D). On the other hand we could not improve the otherwise poor prediction model for RFS (figure 5D, lower panel).

Finally, we used histology analysis to further support the predictive value of myoCAFs. We performed aSMA staining to identify myoCAFs in a tissue micro-array of NMIBC (figure 5E). Manual scoring of the stainings revealed a trend towards lower recurrence-free survival in patients with high myoCAF staining (Log-rank test p-value = 0.0848) (figure 5G). Although the analysis did not reach statistical significance probably due to the low sample size, we provide further evidence to support that myoCAFs are associated to aggressive NMIBC and that they serve as potential biomarkers of recurrence and progression in NMIBC datasets.

## DISCUSION

Several studies have aimed at characterising the TME of NMIBC, with most investigations making use of transcriptomic-based methods (6–8, 29, 30). While very informative intra-study, this type of analysis provides scores that are difficult to compare between studies and are not well-suited for clinical practice. Immunohistological staining offers comparative quantification of cell types, in addition to relevant spatial information, but is limited by the number of markers that can be analysed simultaneously. Therefore, most studies have focused on specific cell types, finding associations to therapy response and prognosis, but failing to provide a wider picture of the NMIBC TME landscape (31–33). Our study is first to provide a reference map of relative proportions of 11 cell types. Comparison with non-pathological tissue allowed us to show cancer-associated changes in the bladder mucosae. The compartment analysis provides a clear picture of non-previously shown changes in NMIBC affecting the microvasculature, as well as increased fibroblasts and MSCs within the non-immune stroma. The differences in the leukocyte compartment are in line with previous publications, with lower presence of T cells but higher of macrophages and NK cells in tumours, which is further exacerbated in MIBC ((30, 34) and supplementary figure 2C). This suggests a tumor promoting role for these cells, as indicated in therapy response studies (32, 35, 36).

Immune checkpoint inhibitors targeting the PD-1/PD-L1 pathway have shown benefits in MIBC. They also constitute a promising new treatment strategy in BCG-unresponsive high-risk NMIBC patients and are under investigation in BCG naïve high risk NMIBC. Despite the good therapeutic results, the use of these targets as prognosis and predictive biomarkers is much more controversial. Some studies have shown a significant association of high PD-L1 expression to bad prognosis (37, 38), to good prognosis (8, 22, 39, 40), or no association (21, 31, 41, 42). Lack of concordance among techniques is unlikely to cause these discrepancies (43). Nonetheless, most studies measure PD-L1 expression in the whole tissue section and/or separating only tumour and immune-infiltrating cells, lacking the required biological resolution. Our results show broad heterogeneity of PD-L1 expression among TME subsets, suggesting most studies lack the granularity required to properly assess value as a good prognosis biomarker. Furthermore, we demonstrate that, while relevant TME cells express important levels of PD-L1, relative cellular abundancy suggests cancer cells may act as a sink for anti-PD-L1 drugs. In addition, as opposed to other cancers (44), PD-L1 expression in NMIBC cells is similar to healthy urothelium. This suggests that immune evasion via this molecule is a late event in BLCA progression.

CAFs have shown mostly a protumorogenic role in BLCA (45), although resident fibroblasts have been shown to restrain tumour development in the early phases in a carcinogen-induced mouse model (46). These evidences suggest a switch in function as fibroblasts differentiate into CAFs. Most studies addressing CAFs in BLCA have focused only on MIBC or mixed cohorts, leaving NMIBC CAFs broadly ignored. We show here that fibroblasts populate NMIBC, especially in high-grade tumours, with no differences between Ta and T1 (data not shown). These results evidence that fibroblast recruitment depend on cancer cell biology rather that level of invasion. Using different approaches, we found that myoCAFs, but no other CAF subsets, associate with bad prognosis in NMIBC patients, as has been reported in MIBC (47, 48). Using immunohistochemistry, Mezheyeuski et al tested CAF-associated markers in BLCA finding they also associated with high grade NMIBC (49). They found fibroblast activation protein (FAP) as the best predictor of poor outcome although not when separating Ta, T1 and T2-4. Interestingly, aSMA was not statistically associated to worse prognosis in the full cohort but it was for T1 tumours, in accordance with our findings. In addition, a different CAF subset, named irCAFs, have been described in a mix cohort of MIBC and NMIBC patients. irCAFs associate with worse overall survival and neoadjuvant and immunotherapy response, including NMIBC, likely through promotion of cancer stemness (12). Future studies should apply more discriminatory panels, and resolve the spatial distribution of cell types, to better characterise CAF subsets in NMIBC.

We also found that CAF-macrophage crosstalk may change over NMIBC progression. Increased interactions in early phases may correlate with co-attraction of both subsets by BLCA cells via CXCL1 (50) and feedback loops via CCL2 and GM-CSF production by CAFs (51). According to our data, myoCAF-macrophage crosstalk is reduced in high-grade NMIBC, but this interaction is recovered in MIBC as described by others. These observations reflect the dynamic nature of the TME as BLCA progresses, likely influenced by cancer cell intrinsic characteristics plus evolution of the inflammatory milieu since it appears tied to tumour grade progression.

Our results support a detrimental role of myoCAFs in NMIBC. Other authors have proposed that bladder CAFs can promote BLCA growth via various factors (52–54), including TGFβ (55–57), a key factor in myoCAF phisiology (58). We show here that CAFs express PD-L1 and that this expression is increased in the myoCAF subset. It would be interesting to measure expression on PD-L1 in other settings where PD-1/PD-L1 inhibitors are being tested, such as BCG-unresponsive patients.

In conclusion, the NMIBC TME is composed by a heterogeneous distribution of cancer, immune and non-immune cells. These cell types exhibit differential features, such as PD-L1 expression, which study may contribute to improve patient management. Our results strongly suggest that myoCAFs play an important role in NMIBC. Future studies, should elucidate how and why myoCAFs associate with worse prognosis in order to find appropriate targets to modulate their function.

## METHODS

### Patients and samples

We recruited 98 patients with BLCA diagnosis between December 2019 and December 2022 at an academic tertiary referral hospital. Patients with muscle-invasive bladder cancer (MIBC), carcinoma in situ and those from whom we could not obtain an acceptable sample were excluded. The study was enriched for high grade, high risk tumours based on the greater clinical need for this patient population. Clinical and demographic data are summarized in supplementary table 1. Due to the limited number of female patients, inherent to the disease, sex was not considered as a biological variable in this analysis. Nonetheless, we did generated two separate analysis for males and female when validating the results using transcriptomic data, and similar findings are reported for both sexes.

### Tissue digestion and antibody staining for flow cytometry

Tissue biopsies were carefully cut into small pieces and digested using collagenase P at 200 μg/ml (Sigma-Aldrich), dispase II 800 μg/ml (thermofisher scientific) and DNase I 100 μg/ml (Sigma-Aldrich) in DMEM at 37°C in a water bath. After 30 min samples were mixed by pipetting up and down and half of the digestion media replenished by fresh until all tissue was digested. Media replacement was repeated two more times with 15 min incubations in between. After erythrocyte lysis, cell suspensions were centrifuged, resuspended in FACS buffer (PBS 2% BSA) and filtered through a 40 μm cell strainer (Corning). Cells were stained with Zombie aqua (Biolegend) in PBS for 20 min at room temperature and then divided into two tubes and used for immunofluorescence staining. Immunofluorescence staining was performed by incubating the cells in PBS containing 1% BSA and 0.01% Sodium azide in the presence of saturating amounts of fluorochrome-conjugated antibodies for 30 minutes at 4°C. Intracellular stainings were done using Cyto-Fast™ Fix/Perm Buffer Set (BioLegend) following manufacturer’s instructions. All antibodies and reagents used can be found in supplementary table 2. Samples were run in a Fortessa X20 flow cytometer (BD Biosciences) at CIEMAT flow cytometry facilities and analysed using the FlowJo software (FlowJo, LLC). Live cells were gated as Zombie aqua negative and by FSC/SSC parameters and doublets discriminated by comparing FSC-A versus FSC-H.

In order to determine the percentage of PD-L1+ cells, all samples were stained with the full panel minus anti-PD-L1 (FMO panel) to stablish the cut-off for positivity. The gate for PD-L1+ was set between 1 and 2% in the FMO panel for each cell compartment and the percentage of PD-L1+ was calculated according to the following formula:

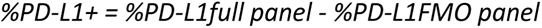

### Computational analysis of flow cytometry data

An optimal compensation matrix was applied to each data set. For automated analysis, we applied a pipeline of analysis for the highest quality data. To avoid any undesirable events or processing artefacts that could potentially affect the downstream results, data were automatically cleaned-up via FlowAI algorithm. Further cleaning was performed by manual gating to remove doublets, debris, and dead cells and the scaling of the data was checked to ensure the arcsinh transformation was unimodal around 0 (values around 150). All files were subsampled to a maximum (equal for all files). Approximately 140,000 target count cells were analysed for each panel. optSNE and FlowSOM were run using default settings, including all markers but zombie aqua and PD-L1. The elbow and consensus metaclustering methods were used and compared, and the optimal number of metaclusters was selected according to results when comparing non-pathological tissue and tumours. EdgeR algorithm was used for all two-group comparisons, using the p-value as the cut-off for statistical significance. Identification of differentially abundant clusters was done based on the expression for all markers (using NxN plot representation), showing concordance between manual and automated analysis techniques. In brief, we used the results from NPT vs tumour comparisons using manual gating as our ground truth. This allowed as to better assess efficient number of metaclusters. The elbow clustering method resulted in excessive division of cellular subsets and chance discovery. We turned to consensus metaclustering since it paralleled more accurately results from the manual gating (supplementary figure 4).

### IHC and immunofluorescence

Nineteen FFPE BLCA sections (7 G1, 9 G2 and 2 G3) were stained with antibodies against aSMA, CD163, PanCK and DAPI. Antigen retrieval was performed using a pressure cooker (Dako, Agilent Technologies). Primary antibodies were incubated overnight. Antibodies used are shown in supplementary table 2. The IHC signal was amplified with a biotin-avidin-peroxidase system (ABC Elite Kit Vector) and visualized using diaminobenzidine (DAB Kit, Vector Laboratories). For immunofluorescence, 4′,6-diamidino-2-phenylindole (DAPI) was used to stain nuclei and images were taken with a Zeiss Axioimager 2 fluorescence microscope. Cytometry analysis of immunofluorescent tissue sections can be found in supplementary methods.

### Cytometry analysis of immunofluorescent tissue sections

Nineteen FFPE BLCA sections (7 G1, 9 G2 and 2 G3) were stained for aSMA, CD163, PanCK and DAPI. Cytometric analysis of 4-parameter immunofluorescences was carried out using the Qupath software (59). To calculate non-vasculature associated aSMA staining area, a pixel classifier was trained with 20+ tissue sections using as annotations aSMA-negative, aSMA-positive and aSMA-vasculature. Cell detection was used for deconvolution of cells using DAPI staining, and positive cell detection was run to classify CD163-positive and CD163-negative cells. The function distance to annotation 2D was applied to calculate the minimum distance of CD163-negative and CD163-positive cells to non-vasculature aSMA-positive structures.

### Single cell RNA-seq analysis

Single cell data from Chen et al. was analysed from raw fastq data. Briefly, reads were pseudoaligned to GRCh38 cDNA sequence assembly using ‘kallisto bus’ command (Kallisto software, default parameters) and sparse matrices were generated bustools program and BUSspaRse R package as described (60). Individual sample matrices loaded with Seurat package were merged in a common object. Data was normalized (sctransform and log normalization for the top 2000 variable genes) and stored as two different slots of the same Seurat object. Linear dimensionality reduction was performed using the first 20 principal components (PC) according to elbow plot visualization. Non-linear dimensional reduction (UMAP) and clustering were performed using these PCs. Cluster stability was visualized in a clustree analysis (resolution set to 0.2). FindAllMarkers was used for cell annotation. Macrophages and fibroblasts were extracted and reanalysed as an independent object. Functional analysis was performed using VISION R package (61) and the resulting signature scores were incorporated as metadata to the Seurat object. Construction of gene-signature scores was done using 2 and 4-fold change differentially expressed genes for each subset or by previously published gene sets, all shown in supplementary table 3.

### Transcriptomic data

The following three independent transcriptomic cohorts were subjected to subset association with recurrence/progression: Robertson et al (6), UROMOL 2021 (7) and de Jong et al (9). Normalized counts for gene expression were downloaded from the publication source and signature scores were calculated by averaging gene expression z-scores within a signature. Patients were stratified according to signature score quartiles as 25% high, average and 25% low.

### Tissue microarray

Cores from formalin-fix paraffin embedded tissue blocks were used to construct tissue microarrays (TMA; 1.5-mm core diameter), with at least two duplicate cores per case (n = 29), using a standard manual method (Beecher Instruments) (62). TMAs were stained with H&E and sections were reviewed to confirm the presence of representative tumour tissue. Demographic, clinical and pathological data is summarised in supplementary table 4.

### Statistical analysis

Categorical variables were expressed as absolute and relative frequency. Continuous variables were expressed as median (interquartile range; IQR) according to a normality test (Kolmogorov– Smirnov test). Comparisons were performed using the Wilcoxon–Mann–Whitney test (for two groups) or the Kruskal–Wallis test (for more than two groups) with Dunńs multiple comparison test. The Spearman correlation analysis method was used to determine the correlation strength and direction between variables.

The survival analyses were performed using the Kaplan–Meier method and described by median and range. Differences between groups were tested using the log-rank test. A Cox proportional hazards model was fitted to estimate hazard ratio (HR) and the corresponding 95% confidence interval (CI). A multivariable model was created with all confounding and relevant factors and had a p-value of <0.1 in the univariate analysis. The best multivariable statistical model was selected using Akaike Information Criterion.

A logistic regression model was created to evaluate the risk factors associated with tumor grade. The study was completed with a multivariable analysis. A step-by-step selection process was used to highlight the most relevant factors, to identify the significant sets among variables and to avoid confusion in the model. We used odds ratios (ORs) and 95% confidence intervals (CIs) to present the results of the regression analysis.

All analyses were done using Stata InterCooled for Windows version 16 (StataCorp. 2019. Stata Statistical Software Release 16, StataCorp LLC, College Station, TX, USA), GraphPad Prism version 9 for Windows (GraphPad Software, Boston, Massachusetts USA) and R (version 4.2.1; R Foundation for Statistical Computing, Vienna, Austria) and a level of significance of 5%.

### Study approval

This study was approved by Research Ethics Committee of the University Hospital “12 de Octubre” and informed consent was obtained from all patients.

### Data availability

Values for all data points in graphs are reported in the Supporting Data Values file.

## Supporting information

Gomez C et al Supplementary materials

Gomez C et al Supplementary tables

## AUTHOR CONTRIBUTIONS

CGC: designing research studies, analyzing data, writing the manuscript, collection of samples and clinical data

FGR: providing reagents and collection of samples and clinical data

MPE: conducting experiments, acquiring data

IL: conducting experiments, acquiring data

CSC: designing research studies

LM: conducting experiments, acquiring data

SPN: conducting experiments, acquiring data

EMM: conducting experiments, acquiring data

CR: conducting experiments, acquiring data

RS: analyzing data

MRI: collection of samples and clinical data

JMV: analyzing data, writing the manuscript

FXR: designing research studies, writing the manuscript

DC: providing reagents

CMA: analyzing data

ARA: collection of samples and clinical data

MD: designing research studies, providing reagents, writing the manuscript

JMP: designing research studies, providing reagents, writing the manuscript

VGM: designing research studies, conducting experiments, acquiring data, analyzing data, providing reagents, writing the manuscript

CGC: designing research studies, analyzing data, writing the manuscript, collection of samples and clinical data

FGR: providing reagents and collection of samples and clinical data MPE: conducting experiments, acquiring data

IL: conducting experiments, acquiring data CSC: designing research studies

LM: conducting experiments, acquiring data SPN: conducting experiments, acquiring data EMM: conducting experiments, acquiring data CR: conducting experiments, acquiring data RS: analyzing data

MRI: collection of samples and clinical data JMV: analyzing data, writing the manuscript

FXR: designing research studies, writing the manuscript DC: providing reagents

CMA: analyzing data

ARA: collection of samples and clinical data

MD: designing research studies, providing reagents, writing the manuscript JMP: designing research studies, providing reagents, writing the manuscript

VGM: designing research studies, conducting experiments, acquiring data, analyzing data, providing reagents, writing the manuscript

## ACKNOWLEDGMENTS

We thank the Histology Laboratory from CIEMAT, namely Pilar Hernandez Lorenzo, for the histological processing of tumour samples and the Laboratory of Cytometry and Cellular Separation, specifically Omaira Alberquilla for their help with the flow cytometry protocols and analyses. We acknowledge Miriam Marques, from the Epithelial Carcinogenesis Group, Spanish National Cancer Centre-CNIO (Madrid) for critical support in manuscript preparation. We acknowledge the following funding bodies:

NCT04134000 clinical trial, funded by Hoffmann-La Roche: Conduct of the study; collection, management of the data.

Grant SAF2015-66015-R and PID2019-110758RB-I00, funded by MCIN/AEI/10.13039/501100011033: design and conduct of the study; collection, management, analysis, and interpretation of the data; and preparation, review, or approval of the manuscript

Grant CIBERONC number CB16/12/00228, funded by Instituto de Salud Carlos III: Design and conduct of the study; collection, management, analysis, and interpretation of the data; and preparation, review, or approval of the manuscript.

V.G.M. supported by fellowships 2018-T2/BMD-10342 funded by Consejería de educación, universidades, ciencia y portavocia de la Comunidad de Madrid, INVES222946GARC funded by Fundación Científica de la Asociación Española Contra el Cáncer: Design and conduct of the study; collection, management, analysis, and interpretation of the data; and preparation, review, or approval of the manuscript.

S.P.N. supported by fellowship SFRH/BD/ 144241/2019) funded by FCT-Fundação para a Ciência e Tecnologia: Conduct of the study; Collection of data; review of the manuscript.

I.L. is supported by a predoctoral fellowship PRDMA19024LODE, funded by Fundación Científica de la Asociación Española Contra el Cáncer: Conduct of the study; Collection of data; review of the manuscript.

L.M. supported by fellowship POSTD19036MORA, funded by Fundación Científica de la Asociación Española Contra el Cáncer: Conduct of the study; Collection of data; review of the manuscript.

