## Supplementary material for "Comprehensive Landscape of Non-muscle Invasive Bladder Cancer Tumour Microenvironment and Prognostic value of Cancer-Associated Myofibroblasts": Gomez C et al Supplementary materials

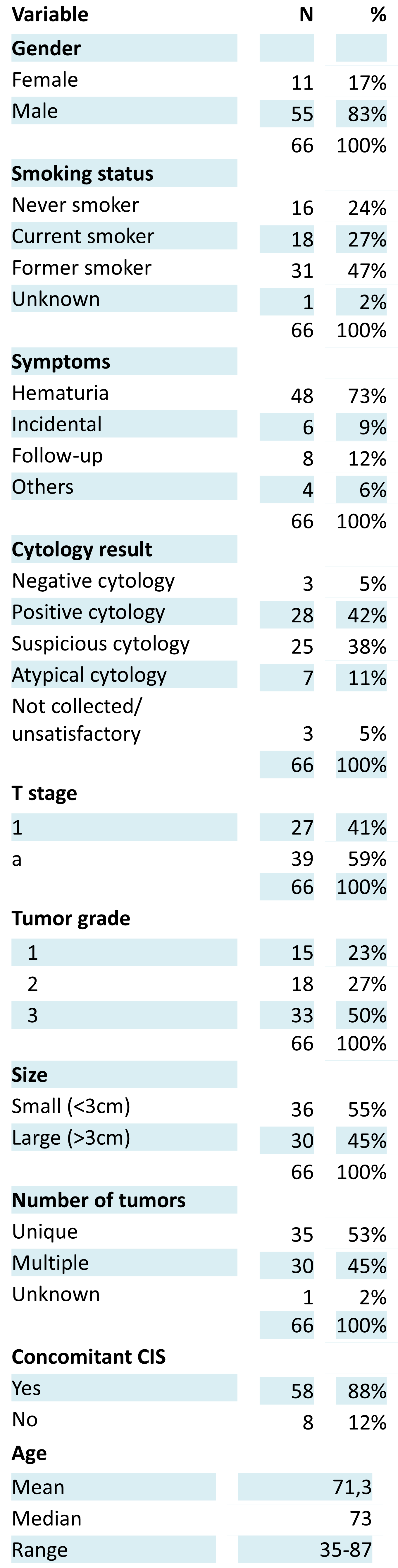

**Supplementary table 1**. Demographic, clinical and pathological data form patients included in the study.

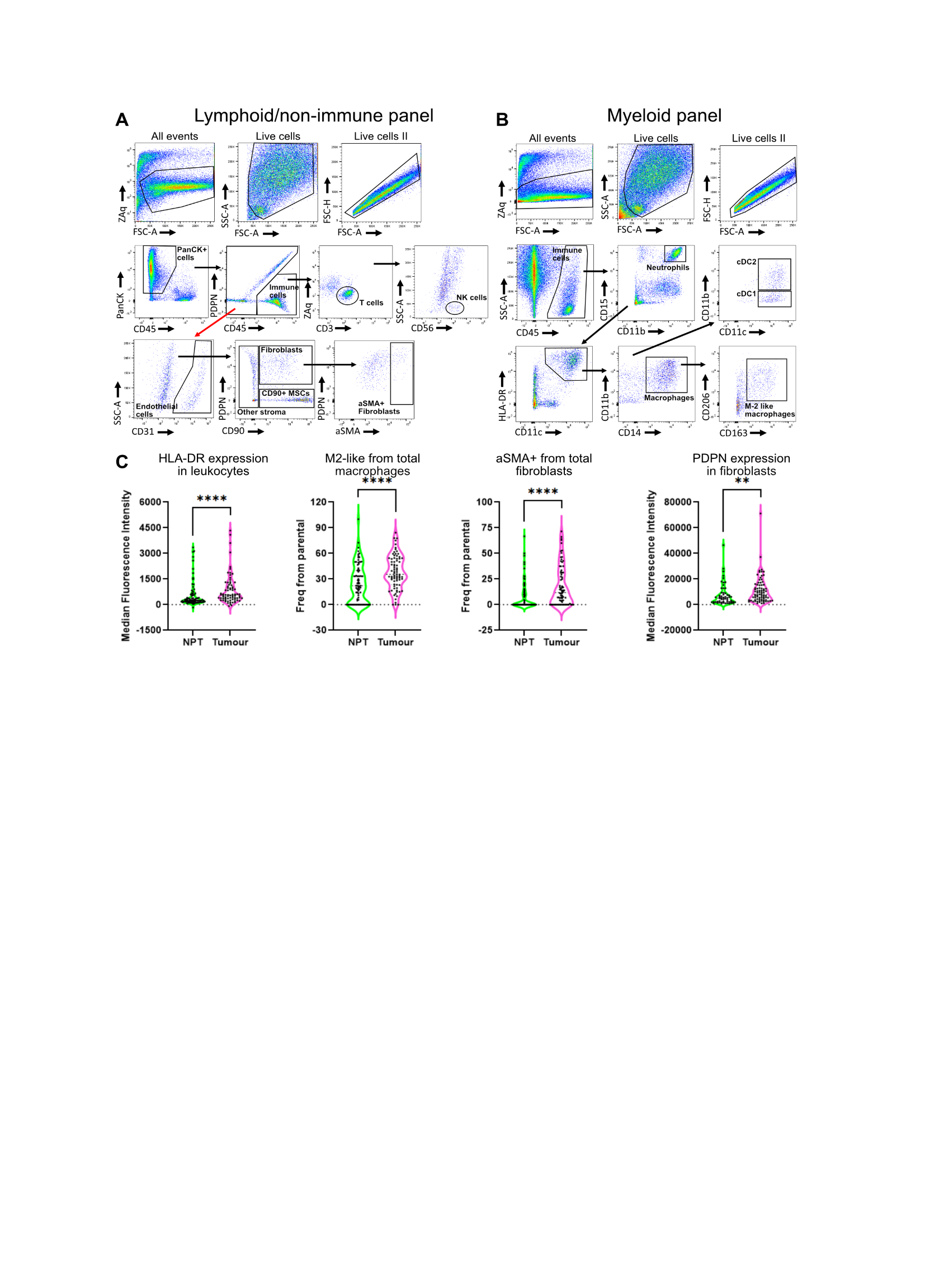

**Supplementary figure 1. NMIBC presents inflammation-associated features**. Conventional gating strategy for the quantification of lymphoid/non-immune (A) and myeloid cells (B) using two separate flow cytometry panels. C) Comparisons between non-pathological tissue (NPT) and tumours for several inflammation surrogate markers. Frequencies from parental gates and median fluorescence intensities are shown. P-values * < 0.05; *** < 0.0005 by Wilcoxon–Mann–Whitney test. HLA-DR, Human Leukocyte Antigen – DR Isotype; PDPN, podoplanin; aSMA, alpha-smooth muscle actin; MSC, mesenchymal stromal cells.

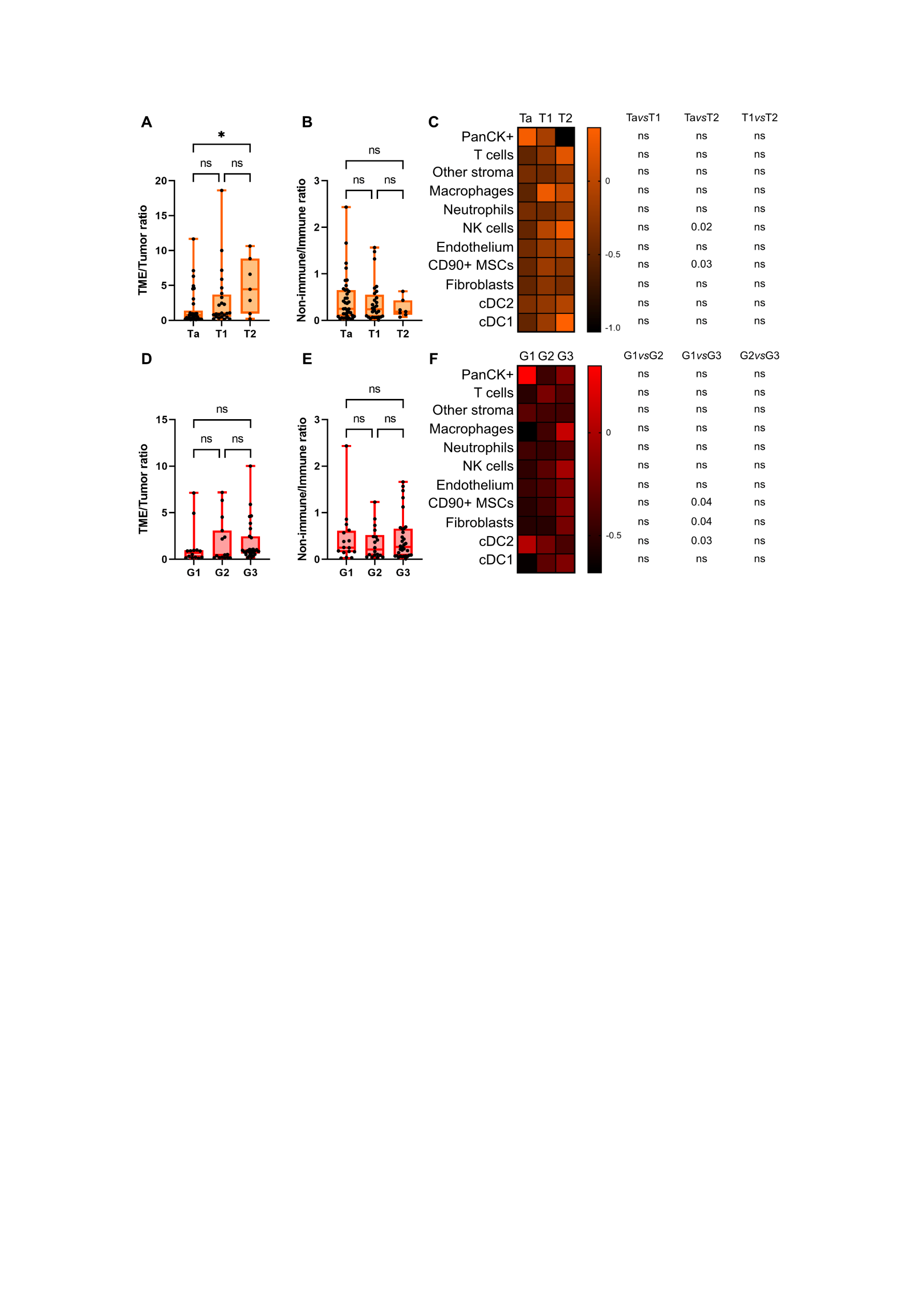

**Supplementary figure 2. Dynamics of tumour microenvironment composition along bladder cancer development.** Flow cytometry data was used to analyse frequency of cell compartments and subtypes in patients stratified by pT stage and grade. For comparisons between pT stages, 6 to 8 T2 tumours were added to the cohort. A, D) Ratios for stroma/cancer cell proportions. B, E) Ratios of non-immune/immune cell proportions were calculated from PanCK-negative cells. C, F) Heatmaps show medians for the proportion from total cells of the indicated cell subtypes along pT stage and grade. P-values are shown for statistically significant comparisons by Kruskal-Wallis test (ns, non-significant) applying Dunn´s test for multiple comparisons.

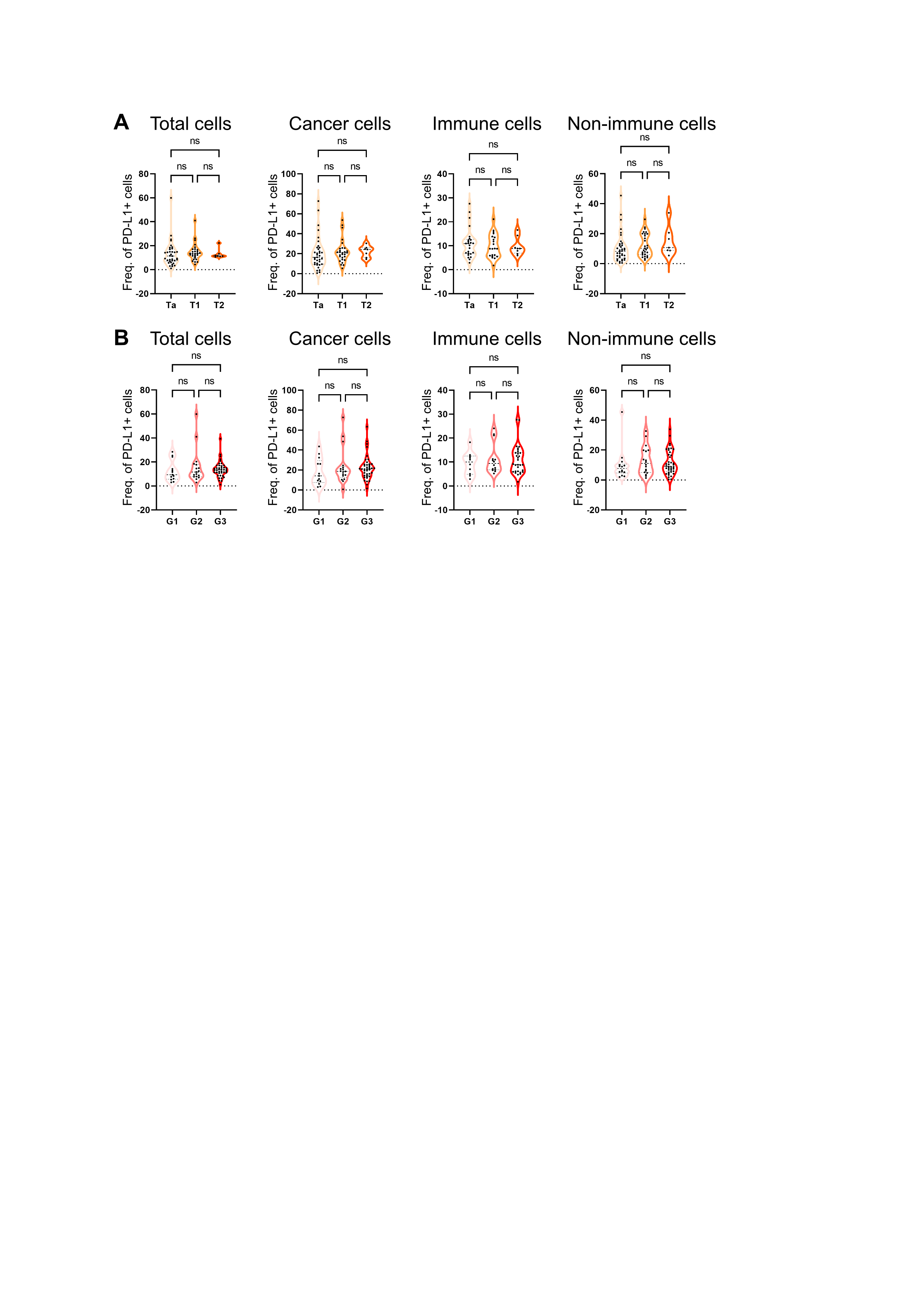

**Supplementary figure 3. No association between PD-L1 expression in different cell compartments and pT stage and grade.** Flow cytometry data was used to calculate the percentage of PD-L1+ cells within the indicated cellular compartments in patients grouped according to pT stage (A) and grade (B). Each dot represents one patient. For pT stage analysis, 6-8 T2 tumours were included. ns, non-significant by Kruskal-Wallis test.

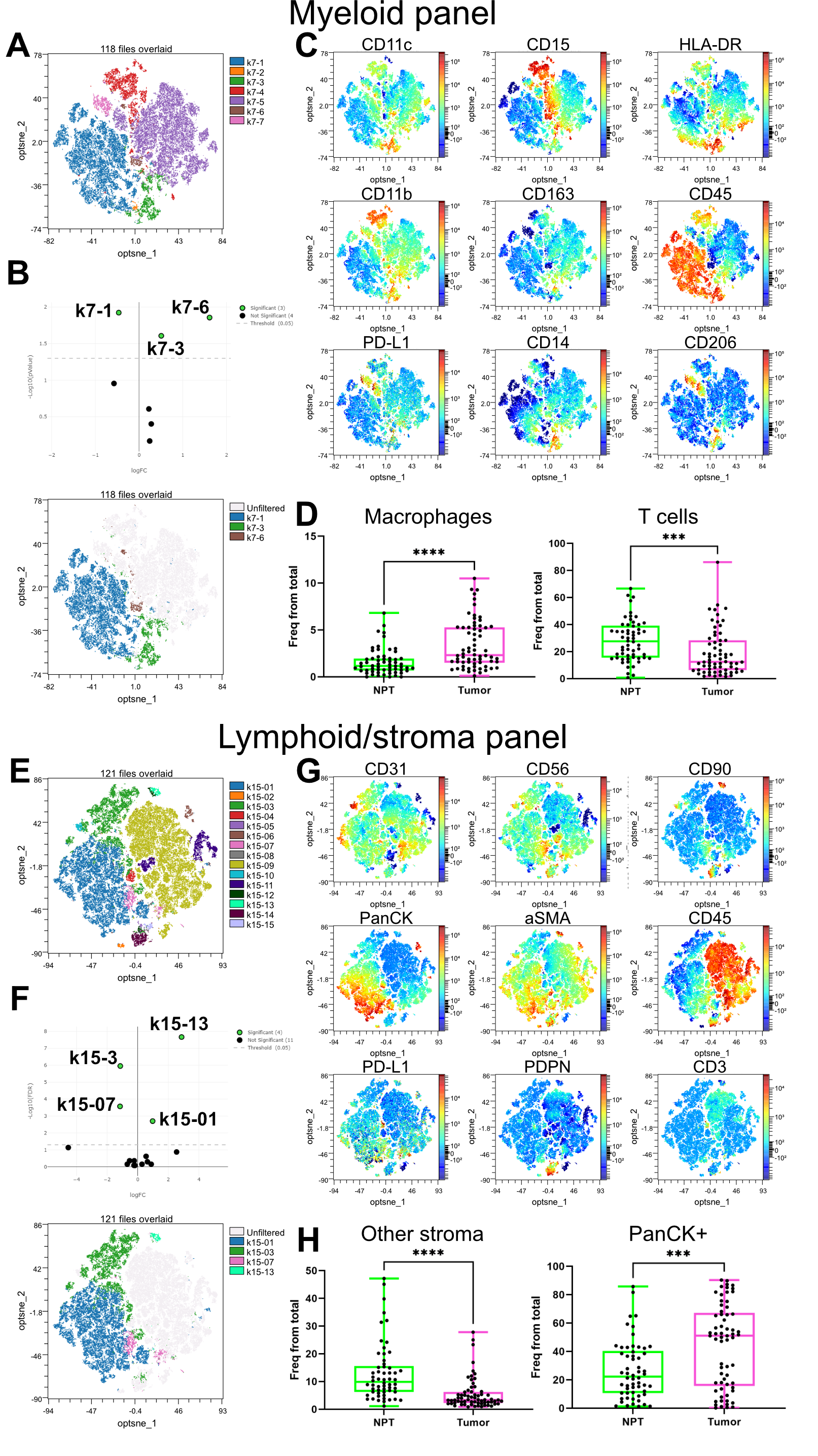

**Supplementary figure 4. Semi-supervised computational analysis of two flow cytometry dataset to compare non-pathological tissue (NPT) and tumours.** Datasets corresponding to the flow cytometry panels for myeloid (A-C) a lymphoid/non-immune stroma cells were subjected to computational analysis using the OMIQ platform. First, reduction of dimensions was performed by optSNE and semi-supervised clustering by FlowSOM, setting the optimal number of clusters as 7 and 15 respectively. A, E) Visualisation of cell clusters overlaid in optSNE maps. B, F) Volcano plots show clusters presenting statistically different frequency in tumours by EdgeR algorithm. optSNE maps highlight over (left side) and under (right side) represented clusters in tumours. C, G) Color-coded optSNE maps showing the expression of the indicated cell markers. D, H) Validation of computational results using the manual gating strategy. D) k7-3 and k7-1 corresponds to macrophages and T cells respectively. H) k15-01 and k15-03 corresponding to cancer cells and other stroma cell subsets respectively. P-values *** < 0.0005 and **** < 0.0001 by Wilcoxon–Mann–Whitney test.

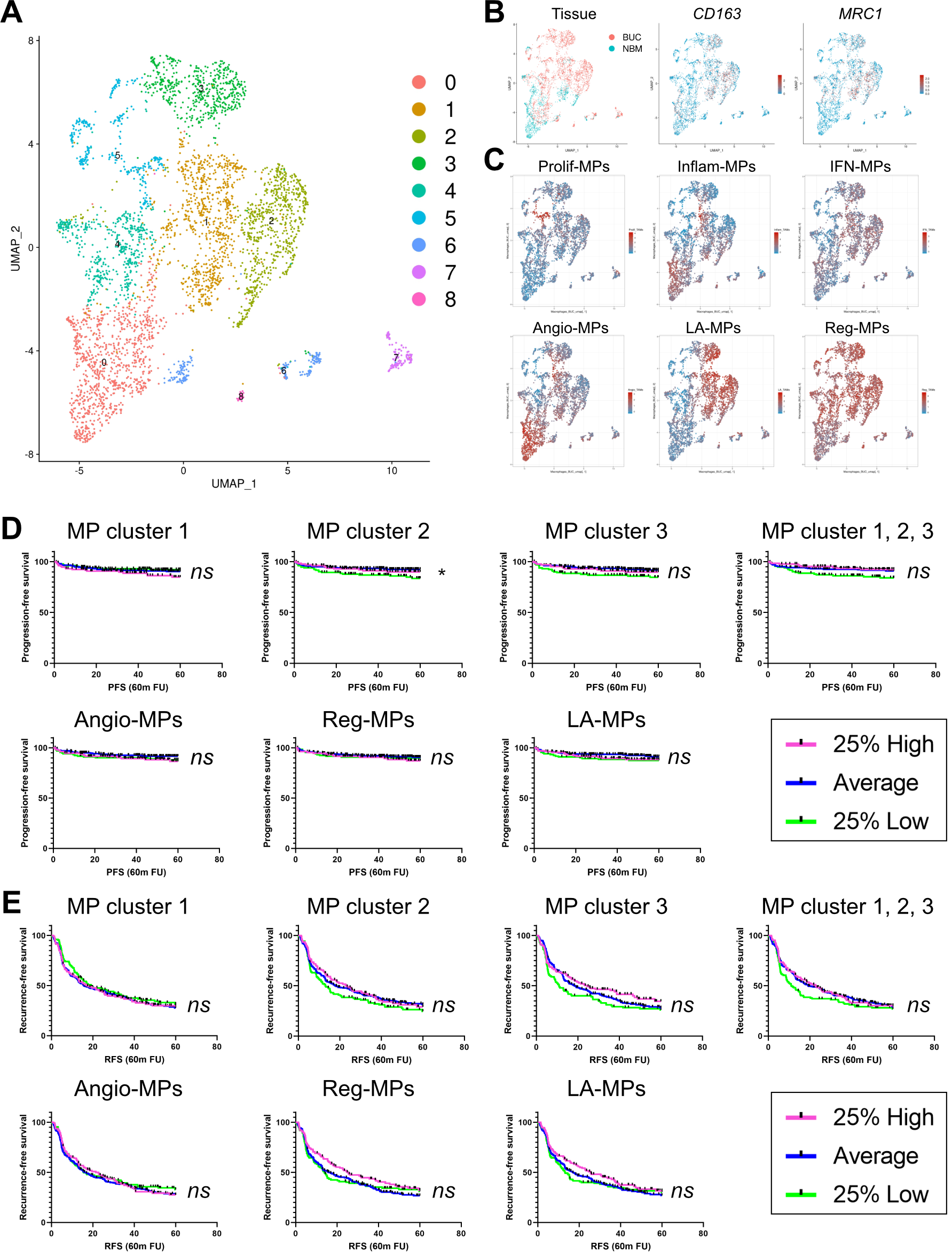

**Supplementary figure 5. Tumour-associated macrophage subsets do not correlate to prognosis in NMIBC**. Single cell data from Chen et al. was analysed from raw fastq data and macrophages (MP) were extracted and reanalysed as an independent object. Functional analysis was performed using VISION R package and the resultant signature scores were incorporated as metadata to the Seurat object. UMAP plots color-coded for macrophage clusters (A), bladder cancer (BC) versus non-pathological tissue (NPT) and CD163 and MRC1 expression (B), gene-signature scores for described tumour-associated macrophage subsets. D-E) CD163/MRC1-expression associated macrophage clusters were selected for further analysis. Kaplan–Meier plots showing probability of recurrence-free and progression free survival of patients stratified according to gen-signature scores for the indicated macrophage subsets. Log-rank (Mantel-Cox) test was used to calculate statistical significance. * p-value < 0.05; ns, not significant. Angio-MP, angiogenic macrophages; LA-MP, lipid associated macrophages; Reg-MP, regulatory macrophages; Prolif-MP, proliferative macrophages; Inflam-MP, inflammatory macrophages; IFN-MP, interferon macrophages. FU, follow-up

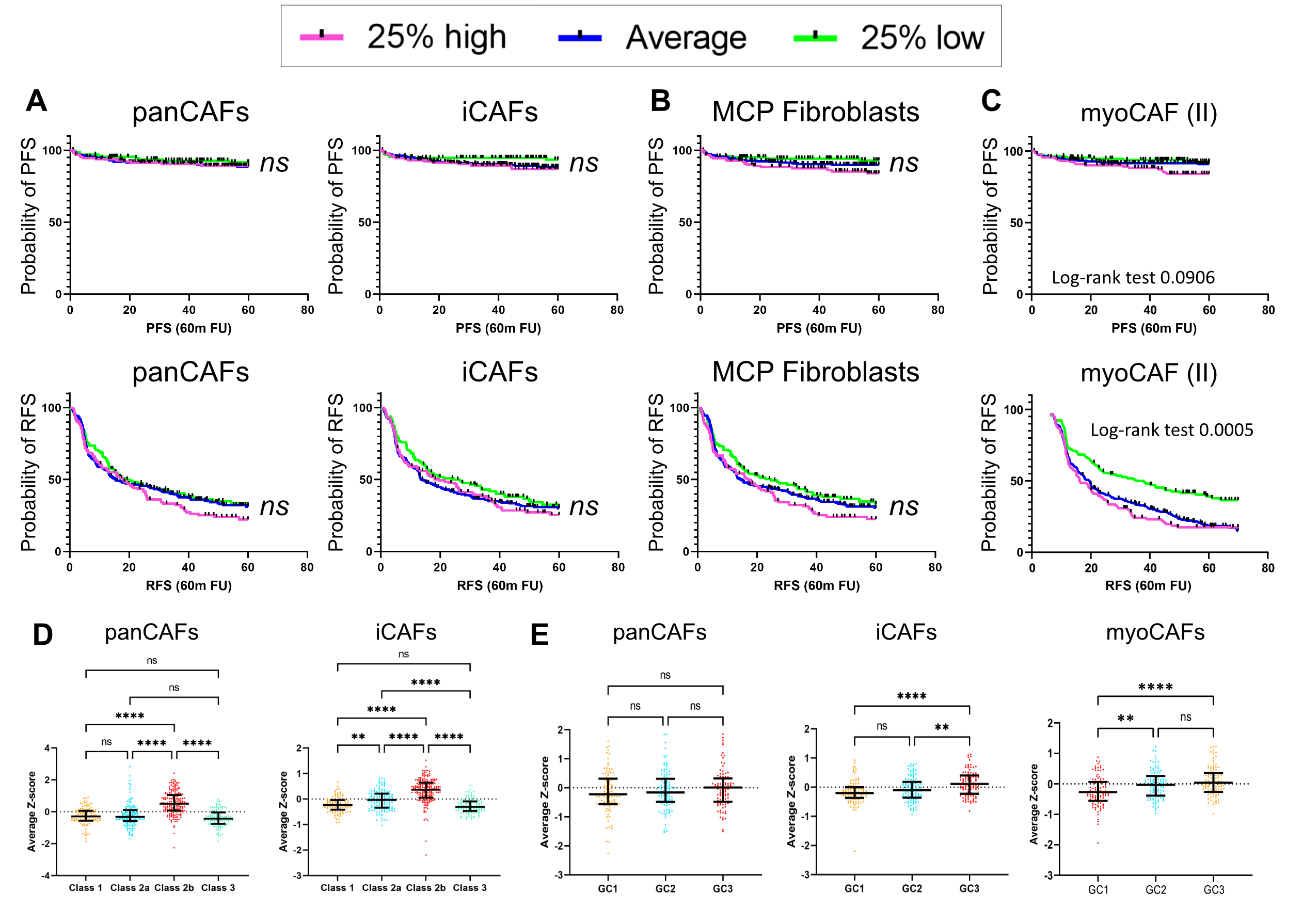

**Supplementary figure 6. Correlation of fibroblast subset gene signatures to prognosis in NMIBC.** Different gene signature scores were generated for CAF subsets to challenge clinical outcome of the UROMOL 2021 cohort of NMIBC tumours. Patients were ranked according to these scores and three groups were formed. A-C) Kaplan–Meier plots for probability of progression-free survival (PFS, upper panel) and recurrence-free survival (RFS, lower panels) for the indicated subsets are shown. Log-rank (Mantel-Cox) test was used to calculate statistical significance between curves. A) Total CAFs (panCAF) and inflammatory CAFs (iCAFs). B) MCP counter fibroblast signature. C) MyoCAF signature newly generated from Chen et al. D-E) panCAF and iCAFs gene-signature scores in patients stratified by UROMOL 2021 transcriptomic classes (D) and genomic classes (E). F) Overview of hazard ratios calculated from univariate Cox regressions of recurrence-free (upper panel) and progression-free (lower panel) survival using clinical and molecular features. Dots indicate hazard ratios and horizontal lines show 95% confidence intervals (CI). P-values and sample sizes, n, used to derive statistics are indicated. CIS, carcinoma in situ; myoCAF, cancer-associated myofibroblasts.

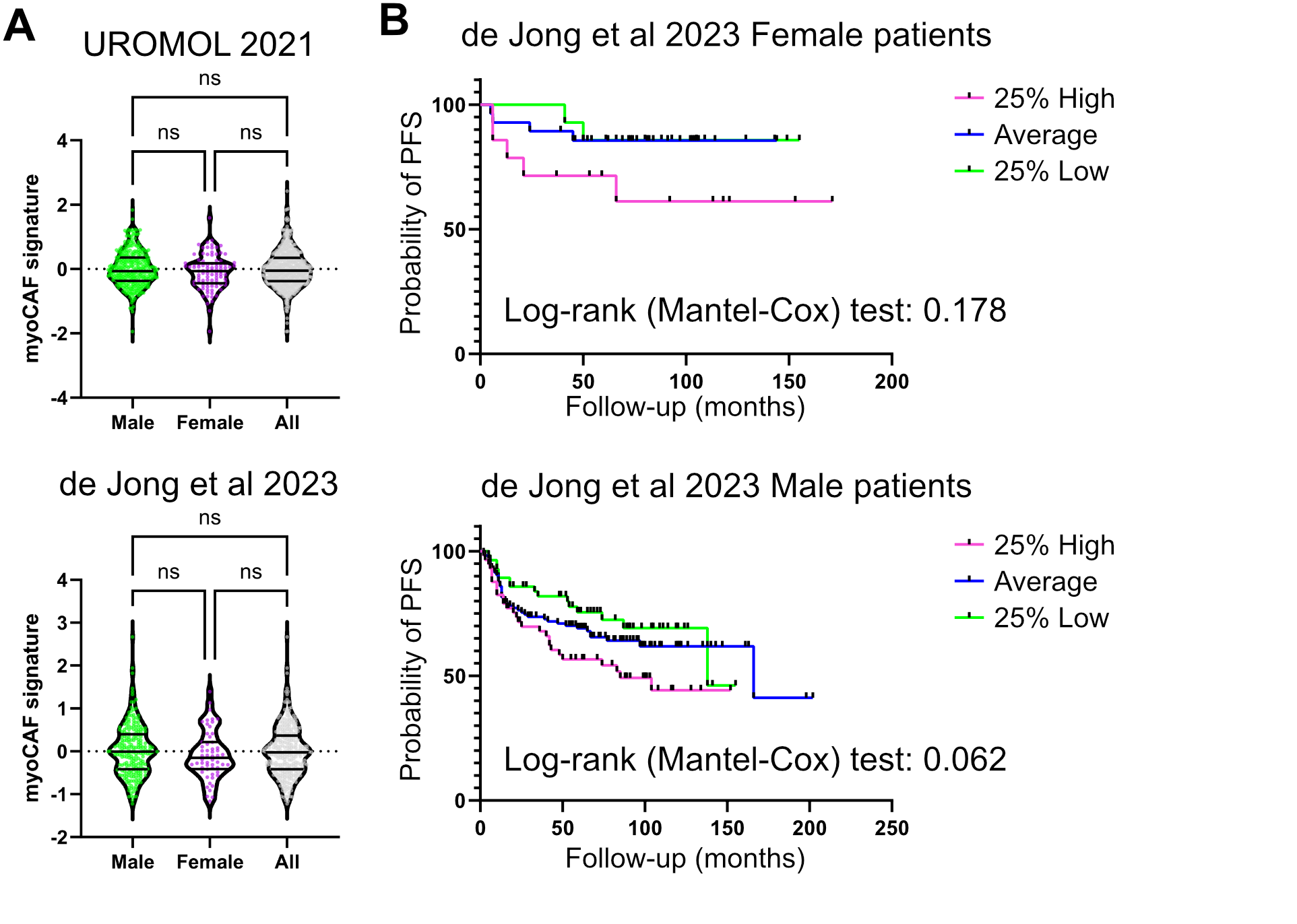

**Supplementary figure 7. myoCAF score shows similar association with poor prognosis in male and female NMIBC patients**. Gene signature scores for myoCAFs were compared between all, male and female patients in the two indicated NMIBC transcriptomic cohorts. Statistical analysis was done with Kruskal-Wallis test with Dunn´s correction for multiple comparisons. ns means non stastistical results. B) Female and male patients were ranked according to myoCAF scores and three groups were formed. Figure shows Kaplan–Meier plots for probability of progression-free survival (PFS) for the indicated subsets. Log-rank (Mantel-Cox) test was used to calculate statistical significance between curves.
